# Molecular and genetic regulation of pig pancreatic islet cell development

**DOI:** 10.1101/717090

**Authors:** Seokho Kim, Robert L. Whitener, Heshan Peiris, Xueying Gu, Charles A. Chang, Jonathan Y. Lam, Joan Camunas-Soler, Insung Park, Romina J. Bevacqua, Krissie Tellez, Stephen R. Quake, Jonathan R. T. Lakey, Rita Bottino, Pablo J. Ross, Seung K. Kim

## Abstract

Reliance on rodents for understanding pancreatic genetics, development and islet function could limit progress in developing interventions for human diseases like diabetes mellitus. Similarities of pancreas morphology and function suggest that porcine and human pancreas developmental biology may have useful homologies. However, little is known about pig pancreas development. To fill this knowledge gap, we investigated fetal and neonatal pig pancreas at multiple, crucial developmental stages using modern experimental approaches. Purification of islet β-, α- and δ-cells followed by transcriptome analysis (RNA-Seq) and immunohistology identified cell- and stage-specific regulation, and revealed that pig and human islet cells share characteristic features not observed in mice. Morphometric analysis also revealed endocrine cell allocation and architectural similarities between pig and human islets. Our analysis unveiled scores of signaling pathways linked to native islet β-cell functional maturation, including evidence of fetal α-cell GLP-1 production and signaling to β-cells. Thus, the findings and resources detailed here show how pig pancreatic islet studies complement other systems for understanding the developmental programs that generate functional islet cells, and that are relevant to human pancreatic diseases.

**Summary Statement:** This study reveals transcriptional, signaling and cellular programs governing pig pancreatic islet development, including striking similarities to human islet ontogeny, providing a novel resource for advancing human islet replacement strategies.

## Introduction

Progress from studies of pancreas biology in humans (Hart and Powers, 2019; Hrvatin et al., 2014b) has advanced our understanding of post-natal islet regulation and function. The recognition that mouse pancreas biology – while essential – has limitations for understanding human pancreas formation or diseases (Hattersley and Patel, 2017; Maestro et al., 2007), has intensified interest in experimental systems that more closely reflect human pancreas development (McKnight et al., 2010; Pan and Brissova, 2014). There is a specific, crucial knowledge gap in our understanding of human islet and pancreas development from mid-gestation through neonatal and juvenile stages, a period when critical aspects of islet development are known to occur (Arda et al., 2016). The use of human primary tissues to address such questions is limited by inter-individual heterogeneity, unpredictable and restricted access to tissue from key developmental stages, and challenges implementing high-throughput molecular approaches. Likewise, reconstitution of islet development from stem cell sources or fetal human cell lines remains imperfect (Sneddon et al., 2018), thereby limiting interpretation of developmental studies in those systems.

Humans and pigs are omnivorous mammals with similar physiology and metabolic diseases, including diabetes mellitus and its complications, from diet-induced obesity, insulinopathy and β- cell stress (Dyson et al., 2006; Lim et al., 2018; Renner et al., 2013). Recent experimental advances expand possible studies of pig pancreas development to include gain- and loss-of-function genetics (Kemter et al., 2017; Matsunari et al., 2013; Sheets et al., 2018; Wu et al., 2017), fetal surgery and immunomodulation (Fisher et al., 2013), and primary islet cell genetics (Peiris et al., 2018). Prior studies of pig pancreas development have largely relied on immunohistological survey of tissue cell types (Carlsson et al., 2010; Ferrer et al., 2008; Hassouna et al., 2018; Nagaya et al., 2019). Moreover, studies linking hallmark islet functions, like regulated insulin secretion and dynamic changes in gene expression at advancing developmental stages, have not been previously reported (Mueller et al., 2013). Thus, pancreas and metabolic research could benefit from systematic application of powerful methods like cell purification by flow cytometry, high-throughput transcriptome analysis, and islet physiological studies across a comprehensive range of pig developmental stages.

Here we apply these and other modern approaches to delineate pig pancreas development, with a focus on islet β-cells, the sole source of insulin, and α-cells, the principal systemic source of the hormone glucagon. Dysregulation of hormone secretion by these cells is thought to be a leading cause of both type 1 diabetes (T1D) and type 2 diabetes (T2D) in humans (Holst et al., 2017; Lee et al., 2016). Our findings provide evidence of greater similarity between pig and human β-cells and α-cells than to cognate mouse cells, and demonstrate how studies of pig pancreatic cells can advance our understanding of the genetic, molecular, signaling and physiological programs that generate functional pancreatic islet cells.

## Results

### Pancreas dissociation and FACS purification of islet cells from fetal and neonatal pigs

To measure gene expression and functional changes in the developing pig pancreas, we established a reliable infrastructure to procure pancreata from fetal or neonatal pigs using rigorous criteria that optimized tissue quality and gene expression analysis (Fig. 1A). To investigate the crucial period of ontogeny culminating in functional islet cells, we systematically isolated pancreata from gestational days (d) 40, 70, 90 and post-natal days (P) 8 and 22 (Fig. 1A). We surveyed sorting methods based on cell surface epitopes, like in prior work with mouse and human cells (Arda et al., 2016; Arda et al., 2018; Sugiyama et al., 2013), but have not yet achieved reliable cell fractionation based with this strategy (R. Whitener and S. K. Kim, unpubl. results). Instead, we used intracellular sorting with specific antibodies against insulin-, glucagon- and somatostatin- to isolate β-, α- and δ-cells, respectively, from fetal or neonatal pancreata (Fig. S1A), like in prior work on human islets (Arda et al., 2016; Hrvatin et al., 2014a; Peiris et al., 2018). Quantitative RT-PCR (qRT-PCR: Fig. 1B) confirmed enrichment or depletion of expected marker genes for β- (*INS*), α- (*GCG*), δ- (*SST*), ductal (*KRT19*) and acinar cells (*AMY2*).

**Figure 1.**
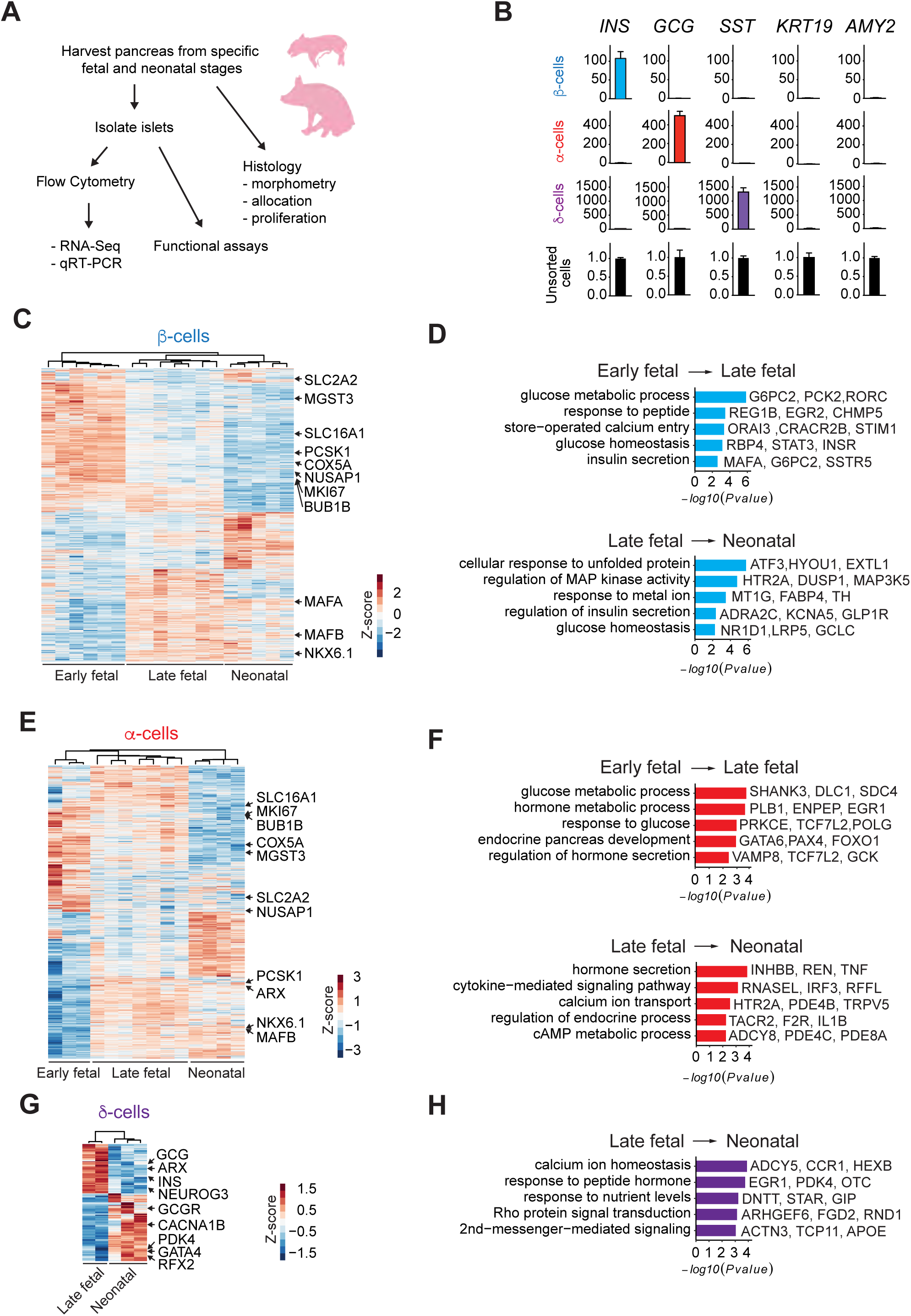
Pancreas dissociation and FACS purification of islet cells from fetal and neonatal pigs. (A) Schematic representation of the study design. (B) qRT-PCR analysis of *INS*, *GCG*, *SST*, *KRT*, and *AMY2* of FACS sorted β-, α- and δ-cell populations from P22 neonatal pigs. The fold change (2^-*ΔΔ*CT) relative to the unsorted cell population is shown on the y-axis. (C) Heatmap of differentially expressed genes (DEGs) in pig β-cells between early fetal, late fetal and neonatal stages. (D) GO Term enrichment analysis of genes increasing with age in pig β-cells. (E) Heatmap of DEGs in pig α-cells between early fetal, late fetal and neonatal stages. (F) GO Term enrichment analysis of genes increasing with age in pig α-cells. (G) Heatmap of DEGs in pig δ-cells between late fetal and neonatal stages. (H) GO Term enrichment analysis of genes increasing with age in pig δ-cells. The Z-score scale represents log_2_(X+1) transformed transcripts per million (TPM) counts. Red and blue color intensity of the Z-score indicates up-regulation and down-regulation, respectively. Adjusted *P* value threshold for GO term analysis was 0.1.

To obtain comprehensive gene expression profiles, we performed RNA-Seq on 37 libraries generated from fetal and neonatal pancreatic cells (Table S1). An average of 7 million reads per sample library uniquely mapped to a current pig reference transcriptome (Sscrofa11.1). Unsupervised hierarchical clustering of transcriptome data with Pearson correlation analysis revealed distinct expression profiles that changed with advancing developmental stage within specific cell types (Fig. S1B). Principal component analysis with these samples showed that transcriptomes clustered in the first principal component according to cell type (β-, α- or δ-cells: PC1, 35% of variance), and according to advancing developmental stage in the second principal component (PC2, 25% of variance: Fig. S1C). Samples from ‘late fetal’ stages d70 and d90 clustered closely and separately from d40 samples, and were grouped together for subsequent analysis (pig development *in utero* averages 114 days). In summary, our cell purification strategy generated high-quality gene expression profiles of developing pig β-, α- and δ-cells.

### High-depth transcriptome maps in pig β-, α- and δ-cells

To evaluate stage-specific differential gene expression, we analyzed transcriptome data in early fetal (d40), late fetal (d70 and d90) and neonatal stages (P8-P22) (Fig. 1C-H). We used the DESeq2 algorithm (Love et al., 2014) to find differentially expressed genes (DEGs) encoding transcripts whose level changed at least 1.5-fold (adjusted *P*<0.05) between these stages in β-, α- or δ-cells. Between early and late fetal stages, we identified 2696 DEGs in β- cells, with transcripts from 1270 genes increased and 1426 decreased. Between late fetal and neonatal stages, we identified 1463 DEGs: transcripts from 785 genes increased, and 678 genes decreased (Fig. 1C and Table S2). In α-cells between early and late fetal stages, we identified 2823 DEGs with transcripts from 1311 genes increased and 1512 genes decreased. Between late fetal and neonatal stages, we identified 2291 DEGs: transcripts from 1200 genes increased, and 1091 genes decreased (Fig. 1E and Table S2). In δ-cells between late fetal and neonatal stages, we identified 1698 DEGs with transcripts from 985 genes increased, and 713 genes decreased (Fig. 1G and Table S2). Among DEGs, we observed subsets of genes enriched in either fetal or neonatal stages in β-, α- or δ-cells, confirming the presence of regulated developmental gene expression programs in specific pig islet cell types. Transcript abundance data from β-, α- and δ-cells are also provided in a searchable dataset (Table S3).

To evaluate coherent patterns of differential gene expression, and effects on signaling pathways at specific stages, we performed gene ontology (GO) term analysis for biological processes with Clusterprofiler (Yu et al., 2012). In the transition from the early to the late fetal stages, we observed enrichment of specific terms linked to islet β- or α-cell development, like glucose homeostasis, endocrine pancreas development, store-operated calcium entry, response to glucose, and regulation of hormone secretion (Fig. 1D,F). In the transition from late fetal and neonatal stages in β-, α- or δ-cells, we observed enrichment of biological processes distinct from those at earlier developmental stages. These included MAP kinase signaling, hormone secretion, cAMP metabolic process, and Rho protein signaling transduction (Fig. 1D,F,H).

### Cell type- and stage-specific gene expression in pig islet β-, α- and δ-cells

Next, we assessed expression of genes thought to regulate development of β-, α- and δ-cells (Fig. 2A-H). Since transcription factors, including PDX1, NKX6.1 and ARX, are known to govern β- and α-cell differentiation in mouse and human islet cells (Aguayo-Mazzucato et al., 2011; Artner et al., 2007; Artner et al., 2010; Collombat et al., 2003; Schaffer et al., 2013), we assessed expression of transcription factors in pigs. Transcripts encoding the factors PDX1, MAFA and NKX6.1 were highly enriched in pig β-cells throughout development, while genes encoding ARX and IRX2 were exclusively expressed in pig α-cells (Fig. 2A, B). Unlike in mice, MAFB is known to be expressed in both human β- and α-cells (Arda et al., 2016; Blodgett et al., 2015). Notably, we observed that *MAFB* was expressed in both pig β- and α-cells (Fig. 2A). As expected, pig δ-cells expressed genes encoding transcription factors such as *HHEX*, *PDX1*, and the transporter *RBP4*, consistent with previous findings in other systems (Muraro et al., 2016) (Fig. 2A,C). GLP1R is known to be highly expressed in human β-cells and involved in β-cell function (Dai et al., 2017). Similar to human, *GLP1R* mRNA was highly expressed in pig β-cells (Fig. 2D). Urocortin3 (UCN3) secreted by β-cells binds to corticotropin-releasing hormone receptor 2 (CRHR2) in δ-cells, stimulating somatostatin secretion (van der Meulen et al., 2015). In accordance with these previous findings, we observed that *UCN3* and *CRHR2* mRNA are highly expressed in pig β-cells and δ-cells, respectively, raising the possibility that this signaling axis is conserved in the pig (Fig. 2D).

**Figure 2.**
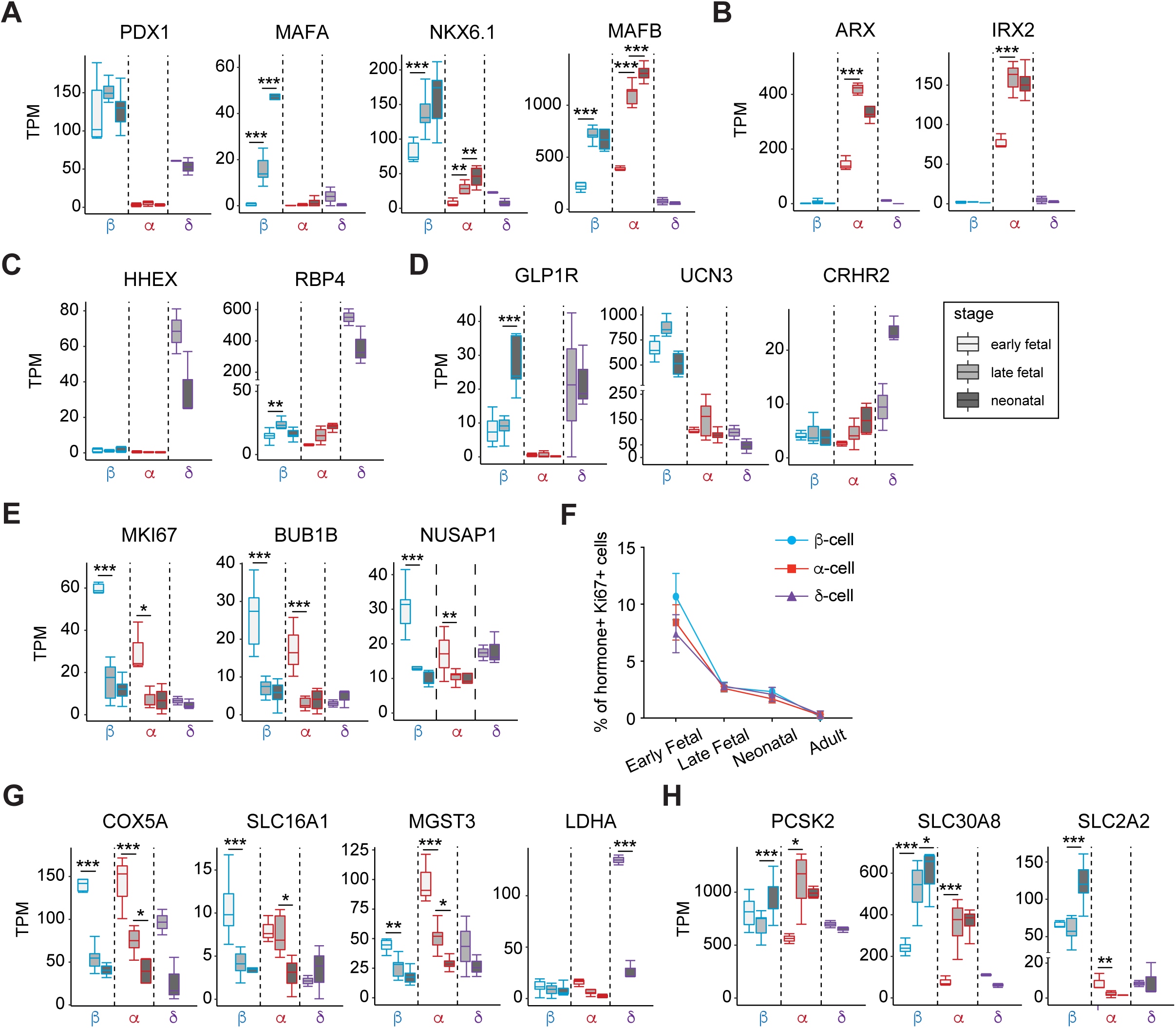
Analysis of gene expression in developing pig islet β-, α- and δ-cells. (A-E,G,H) Boxplots displaying normalized TPM (transcripts per million) counts of β-, α-, and δ-cell specific genes (A-D), markers associated with proliferation (E), disallowed genes (G) and β-cell functional regulators (G). (F) Quantification showing average percentage of INS^+^, GCG^+^ or SST^+^ cells that are Ki67^+^ in each developmental stage (n = 40 images per group, from 3 pigs per group; bars show SD). Adjusted *P* value * *P*≤0.05; ** *P*≤0.01; *** *P*≤0.001

We also observed dynamic expression of genes regulating islet cell proliferation and function in developing pig β- and α-cells (Fig. 2E-H). For example, *MKI67*, *BUB1B* and *NUSAP1*, which encode products known to regulate mitosis and cell cycle progression, declined as β- and α-cell development advanced (Fig. 2E). To validate these findings, we used immunohistology to quantify the nuclear factor Ki67, a marker of cell proliferation encoded by *MKI67* in multiple species, including pigs, humans and mice (Arda et al., 2016; Georgia and Bhushan, 2004; Meier et al., 2008; Teta et al., 2007). In pigs, pancreas immunostaining revealed a high percentage (10%) of Ki67^+^ early fetal islet cells (d40), that declined to 2% by late fetal and neonatal stages, then further declined to <0.1% by adulthood (Fig. 2F and Fig. S2). These changes were mirrored by mRNA levels encoding *MKI67* (Fig. 2E). Thus, we observed evidence of vigorous fetal-stage proliferation of islet cells that, like in mice and humans, declined further after birth.

Likewise, we found that expression of ‘disallowed’ genes like *COX5A, SLC16A1*, *MGST3* and *LDHA*, originally identified in β-cells and thought to restrain β-cell function and maturation (Lemaire et al., 2016; Pullen et al., 2017; Pullen et al., 2010; Sekine et al., 1994), declined during development (Fig. 2G). We also detected dynamic or cell type-specific expression of genes encoding established effectors of islet hormone secretion, like the proprotein convertase *PCSK2*, the ion transporter *SLC30A8*, and the glucose transporter *SLC2A2* (Fig. 2H). These data delineate gene expression changes at genome-scale that orchestrate age-dependent changes in proliferation and maturation of islet β-, α- and δ-cells. Thus, our studies revealed both similarities of (1) *cell type-specific* gene expression between pig and human islet β-, α- and δ-cells, and (2) conserved *stage-specific* gene expression dynamics in pigs.

### Stage-specific islet cell allocation and production of transcription factors

To verify further our RNA-Seq findings in developing islets, we performed immunostaining of pig pancreata at specific developmental stages. At fetal stages, islet β-, α- and δ-cells formed small clusters, which enlarged subsequently in neonatal and adult pancreas (Fig. 3A-C and Fig. S3A). Thus, like in other species (Carlsson et al., 2010; Jeon et al., 2009; Jorgensen et al., 2007), islet cells formed clusters at the inception of their ontogeny, then continued their morphogenesis in multicellular structures through post-natal stages. In fetal, neonatal and adult islets, standard morphometry revealed that differentiation of islet cells into specific lineages was dynamic (Fig. S3A,B). In early fetal pancreas, the number of α-cells is slightly higher than that of β-cells, while the δ-cell fraction was very low (<4%). However, from late fetal to neonatal stages, the number of β-cells is higher than that of α-cells, while the number of δ-cells is similar to that of α-cells (Fig. S3B). Consistent with our RNA-Seq data, antibodies that detect the transcription factors (TFs) PDX1, NKX6.1 and MAFB showed restricted production in specific islet cell subsets. In β-cells, we detected nuclear PDX1, NKX6.1. By contrast, we detected MAFB protein in both β- and α-cells (Fig. 3D-F). Unlike in mice, human β-cells express the transcription factors SIX3 and SIX2 in an age-dependent manner; moreover human α-cells do not express *SIX3* or *SIX2* (Arda et al., 2016). Immunohistology (Fig. 3G) demonstrated SIX3 production specifically in a subset of adult β-cells, and qRT-PCR (Fig. 3H) of pig islets revealed adult-specific production of SIX3. We observed a similar age-dependent expression of *SIX2* (Fig. 3I); the absence of antibodies for pig SIX2 precluded immunohistology studies. In summary, stage-specific and cell-specific expression of multiple TFs in pig islets resembled that observed previously in human islet cell subsets.

**Figure 3.**
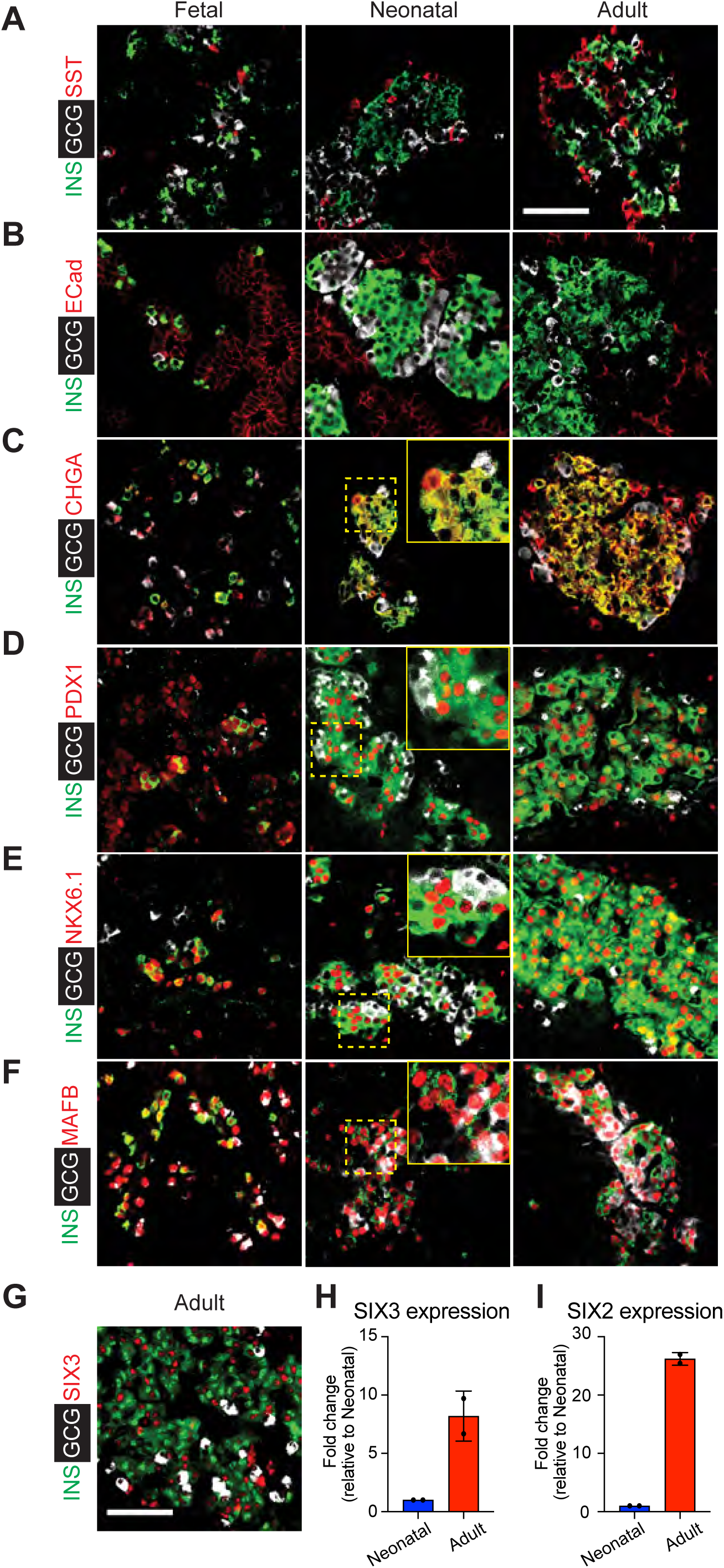
Stage-specific expression of islet factors during development. (A-C) Immunostaining of INS (green) and GCG (white) paired with Somatostatin (SST; red) (A), E-Cadherin (ECad; white) (B), and Chromogranin-A (CHGA; white) (C) in the early fetal, neonatal, and adult pig pancreas. (D-F) Immunostaining of the transcription factors PDX1 (D), NKX6.1 (E), and MAFB (F) in the early fetal, neonatal, and adult pig pancreas. (**G**) Immunostaining of SIX3 with INS and GCG in adult pig pancreas. Insets indicated by the dashed yellow box. Scale bars, 50 µm. (**H, I**) qRT-PCR measures of relative mRNA levels encoding *SIX3* **(H)** and *SIX2* **(I)** in isolated neonatal and adult pig islets (n=2, t-test; Error bars indicate SD). ΔCT values for neonatal and adult *SIX3* expression were 5.1 and 2.0, respectively (compared to β−Actin). ΔCT values for neonatal and adult *SIX2* (compared to β−Actin) were 7.2 and 2.6, respectively.

### Similarities of pig and human β-cells and α-cells

Beyond assessment of TFs, we sought independent evidence of similarities between pig, human and mouse β-cells. We used unsupervised hierarchical clustering with distance matrix analysis of our RNA-Seq data, and extant human and mouse islet cell bulk RNA-Seq data (Arda et al., 2016; Blodgett et al., 2015; Qiu et al., 2017). This analysis suggested that neonatal β-cell transcriptomes in pig and human were globally more similar than to the β-cell transcriptome of mice (Fig. 4A). The absence of sufficient fetal or neonatal mouse α- or δ-cell transcriptome data precluded analogous global comparisons of human, pig and mouse α-cells or δ-cells. GO term enrichment analysis also revealed differential molecular signatures between neonatal pig and juvenile human β-cells. Significantly enriched pig pathways were associated with proliferation, while pathways enriched in human juvenile islet β-cells included those regulating β-cell secretory function (Fig S4 and Table S5). This is consistent with prior observations, including relatively higher proliferation in neonatal pig β-cells compared to juvenile human β-cells (Arda et al., 2016).

**Figure 4.**
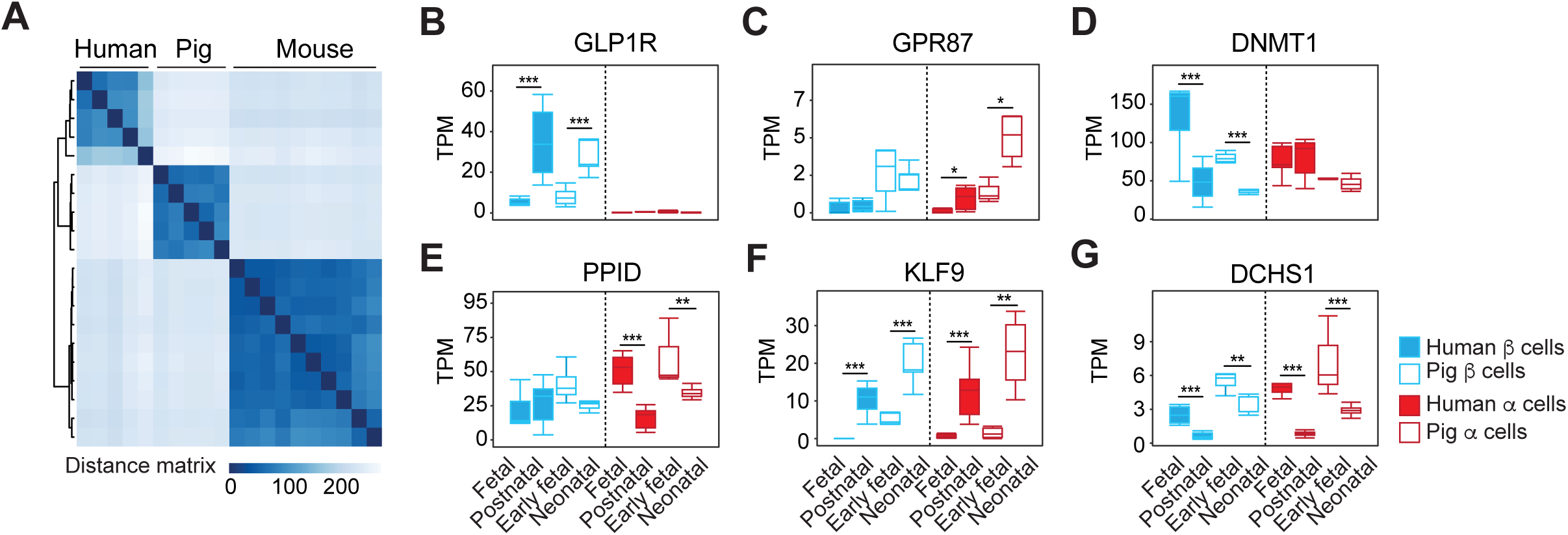
Comparison of gene expression in pig and human β- and α-cell development. (A) Unsupervised hierarchical clustering using distance matrices of neonatal pig, human, and mouse β-cells. (B-G) Boxplots displaying normalized TPM (transcripts per million) counts of select genes whose expression changes in a conserved manner between human and pig development. Adjusted *P* value * *P*≤0.05; ** *P*≤0.01; *** *P*≤0.001.

Next, we performed pairwise comparison of pig and human (Blodgett et al., 2015) cells from fetal and postnatal stages (Fig. 4B-G and Table S4). This analysis revealed conserved cell type-specific and stage-specific regulation of multiple factors. For example, *GLP1R* encodes a G protein-coupled receptor for the incretin hormone GLP-1. Like in humans (Dai et al., 2017), *GLP1R* is expressed in pig β-cells but not α-cells and this expression increases from fetal to neonatal stages (Fig. 4B). By contrast the G protein coupled receptor encoded by *GPR87* is expressed in pig and human α-cells but not β-cells and has increasing expression during postnatal islet maturation (Fig. 4C). Similarities in cell type-specificity or stage-specific expression for other regulators, including *DNMT1*, *PPID*, *KLF9*, and *DCHS1,* provide further evidence of conserved genetic programs underlying pig and human β- and α-cell development (Fig. 4D-G). Taken together, our analysis of specific islet TF’s, global gene expression similarities in β-cells, similarities of developmental islet cell allocation, and of islet morphology provide evidence of remarkable similarities in pig and human β-cell development.

### Dynamic gene regulation during pig β- and α-cell development

Our studies of pig islet development provided an unprecedented opportunity to assess gene expression dynamics throughout fetal and postnatal stages. From the early fetal to neonatal stage, we identified 3,111 genes in β-cells and 3,668 genes in α-cells with dynamic expression (Fig. 5A,B and Table S6). Thus, our analysis permitted grouping of genes in ‘clusters’, based on the sequence of changes observed between three discrete developmental periods. For example, we observed genes in β- and α-cells that increased through late fetal stages, then did <u>n</u>ot <u>c</u>hange thereafter (cluster 2, ‘Up_NC’) and other genes whose transcript levels declined through late fetal stages then did not change thereafter (cluster 7, ‘Down_NC’) (Table S6). We grouped genes in 8 clusters (Up_Down, Down_Up, etc.), and GO term analysis of each cluster revealed dynamic biological processes (Fig. 5A,B and Fig. S5A,B). For example, we observed that the β- and α-cell “Down_NC” clusters were significantly enriched for genes that regulate cell proliferation and cell cycle progression (like *CDK2, CDK14* and *CDC20*; Fig. 5A,B and Table S6). The “Up_NC” cluster in both cell types was enriched for genes associated with multiple signaling pathways involved in glucose and steroid hormone signaling (Fig. S5A,B). Thus, the data identified multiple biological processes involved in islet differentiation and maturation that are dynamically regulated in development.

**Figure 5.**
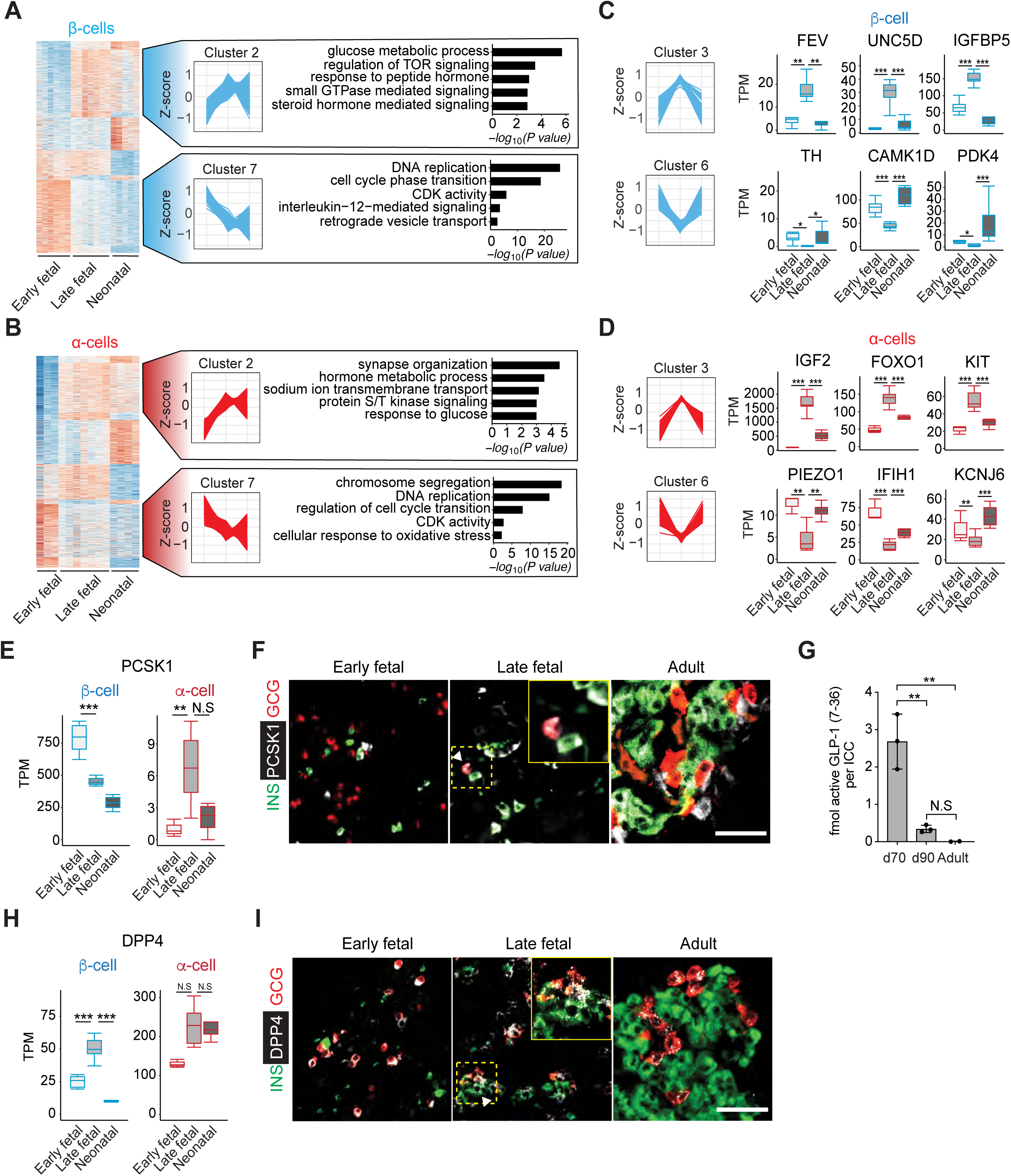
Dynamic gene regulation during pig β- and α-cell development. (A-B) Heat maps of genes with altered expression from early fetal to neonatal stages in β-cells (A) and α-cells (B), showing 8 distinct clusters. Cluster 2 includes genes whose mRNA increased up to late fetal stages then did not change thereafter. Cluster 7 includes genes whose mRNA declined then did not change thereafter. Enriched GO terms in each cluster are shown to the right. Adjusted *P* value threshold was 0.1. The Z-score scale represents log2(X+1) transformed transcripts per million (TPM) counts. Red and blue color intensity of the Z-score indicates up-regulation and down-regulation, respectively. (C-D) The Z-score scale represents log2(X+1) transformed TPM counts with total genes in β-cells (C) and α-cells (D). Cluster 3 (Up-Down) or Cluster 6 (Down_Up). Boxplots displaying normalized TPM (transcripts per million) counts of genes shown. (E) Boxplots displaying normalized TPM (transcripts per million) counts of PCSK1 mRNA. (F) Immunostaining of PCSK1 (white) with insulin (INS; green) and glucagon (GCG; red) in early fetal, late fetal and adult pig pancreas. Insets indicated by the dashed yellow box. White arrows indicate PCSK1^+^ GCG^+^ (double positive) cells. (G) Concentrations of active GLP-1 in lysates from pig islet cell clusters at different developmental stages (ANOVA, *\*\* *P*≤0.01*; bars indicate SD). (H) Boxplots displaying normalized TPM (transcripts per million) counts of *DPP4* mRNA. (I) Immunostaining of DPP4 (white) with insulin (INS; green) and glucagon (GCG; red) in early fetal, neonatal and adult pig pancreas. Insets indicated by the dashed yellow box. White arrows indicate DPP4^+^ INS^+^ (double positive) cells. Adjusted *P* value * *P*≤0.05; ** *P*≤0.01; *** *P*≤0.001, Scale bars, 50 µm.

The frequency of transcript measurement in our workflow also identified unanticipated gene expression dynamics in developing β- and α-cells, particularly in gene clusters with “Up_Down” or “Down_Up” trajectories (clusters 3 and 6: Fig. 5C,D). For example, from fetal to neonatal stages in islet β-cells, we observed increased levels of transcript encoding pyruvate dehydrogenase kinase 4 (PDK4), a factor previously characterized (Pullen et al., 2017) as ‘disallowed’ in β-cells (Fig. 5C). Consistent with this, we observed that expression of *PPARGC1A*, a known positive regulator of *PDK4* transcription (Wende et al., 2005) also increased at this stage in β-cell development (“NC_Up” cluster; Table S6). Thus, unlike other disallowed genes whose expression continuously declined in developing β- and α-cells, *PDK4* expression appeared to increase after late fetal stages, raising the possibility of as-yet unidentified functions for PDK4 in pig β-cell development.

In human diabetes, genetic risk may reflect monogenic or polygenic mechanisms (Cerolsaletti et al., 2019; Hattersley and Patel, 2017; Pearson, 2019; Sanyoura et al., 2018). We hypothesized that causal genes in monogenic forms of diabetes including neonatal diabetes mellitus (NDM) and maturity onset diabetes of the young (MODY) might be developmentally regulated. Thus, we compared expression dynamics of 33 genes, linked by prior studies to human NDM and MODY (Velayos et al., 2017; Yang and Chan, 2016), and with known pig orthologues (Table S7). We found ten genes dynamically expressed between fetal and neonatal stages in pig β-cells. Of those, six genes - *INS, STAT3, NEUROG3, GATA4*, *SLC2A2*, and *PCBD1* - are known to have dynamic expression in human β-cells. In pig α-cells, 13 causal genes are dynamically expressed, with four - *WFS1, NEUROG3, CEL*, and *GATA6* - known to have changing expression in human α-cells. We also identified seven genes which are differentially expressed between the late fetal and neonatal stages in pig δ-cells. Thus our work reveals pig orthologues of causal genes for MODY and NDM whose expression changes in development of multiple islet cell types, identifying opportunities for investigating native dynamic regulation of these genes.

*PCSK1* encodes an endopeptidase normally expressed in intestinal L-cells to regulate GLP-1 production (Rouille et al., 1995; Steiner, 1998), and in β-cells for proinsulin processing. Unexpectedly, we observed a transient increase of *PCSK1* mRNA in fetal α-cells (Fig. 5E); immunostaining confirmed that PCSK1 protein was produced in a subset of late fetal d70 α-cells but not in early fetal or adult α-cells (Fig. 5F). Consistent with these findings, ELISA quantification of bioactive GLP-1 revealed abundant production in d70 islets that decreased thereafter and was undetectable in adult islets (Fig. 5G). Likewise, *DPP4* encodes a protease that inactivates GLP-1 and whose expression is extinguished in postnatal human β-cells (Arda et al., 2016; Blodgett et al., 2015). However, we observed a significant *increase* of *DPP4* mRNA and protein levels in β-cells at late fetal stages, when β-cells also express *GLP1R* (Fig. 5H,I and Table S3). Together, these findings provide evidence that fetal α-cells transiently produce GLP-1, and that GLP-1/GLP1R/DPP4 signaling interactions could regulate fetal β-cell development.

### Genetic regulation of β-cell functional maturation

Age-dependent enhancement of glucose-regulated insulin secretion has been described in rodents and humans (Aguayo-Mazzucato et al., 2011; Arda et al., 2016; Avrahami et al., 2015; Blum et al., 2012; Rorsman et al., 1989). In pigs, prior studies of islet insulin secretion have focused on postnatal stages (Mueller et al., 2013), but did not compare fetal to neonatal stages. We compared glucose-stimulated insulin secretion in islets isolated from late fetal (d70) or weaning-age donors (P22). Glucose or IBMX, a potentiator of intracellular cAMP levels and insulin secretion, did not significantly increase insulin output by fetal islets in static batch cultures (Fig. 6A). Moreover, insulin secretion by fetal islets in the ‘basal’ 2.8 mM glucose condition was higher than that observed from P22 islets (Fig. 6A). In P22 pig islets, glucose or glucose + IBMX significantly increased insulin secretion (Fig. 6A), though to a lesser degree than in human juvenile islets (Arda et al 2016). The pattern of declining basal secretion accompanied by enhanced glucose-regulated insulin secretion in pig islets is similar to findings from prior studies of isolated late-fetal and neonatal rodent islets (Sodoyez-Goffaux et al., 1979), and provides evidence of post-natal pig β-cell functional maturation.

**Figure 6.**
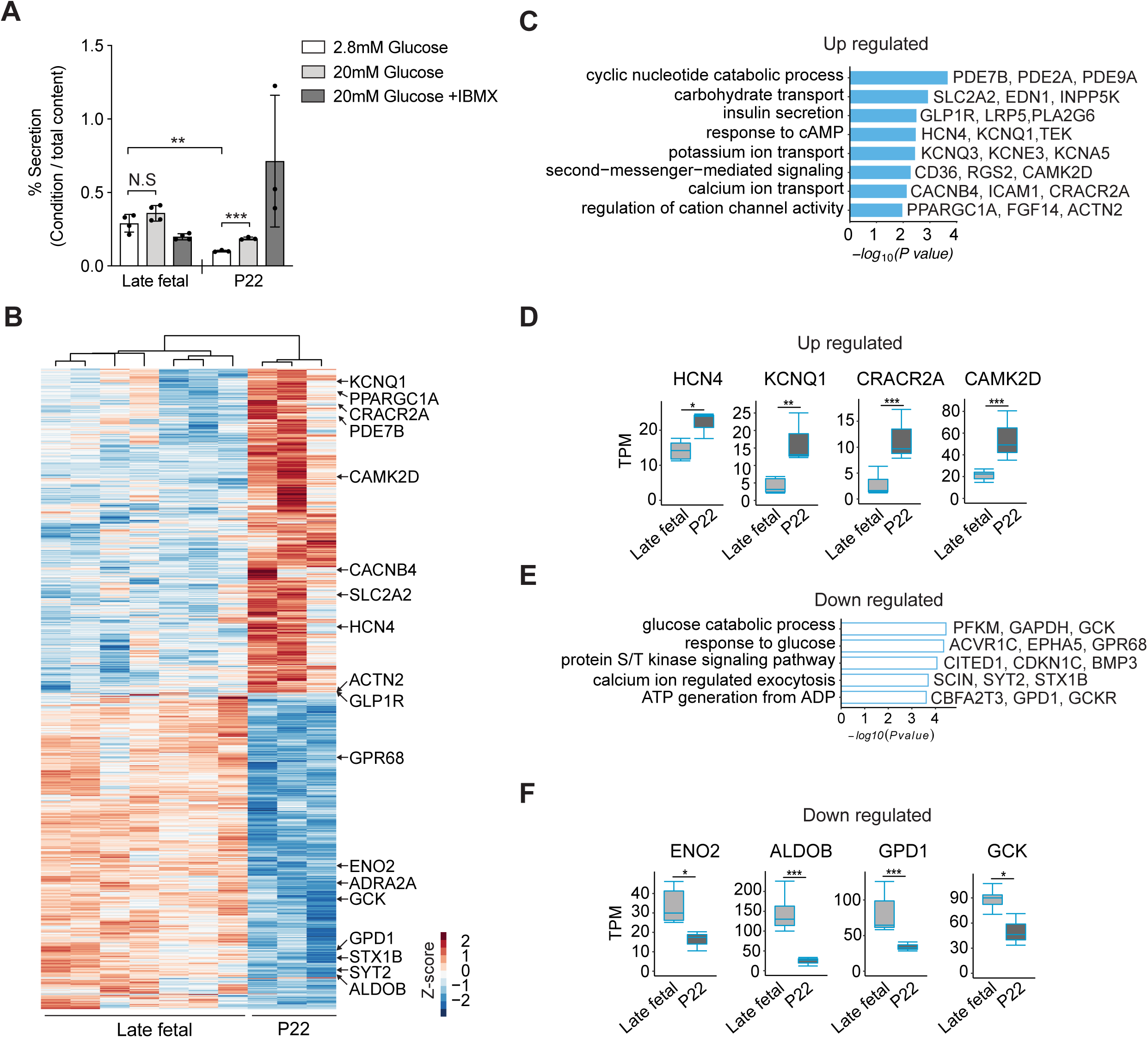
Maturation of β-cell function in postnatal pigs. (A) Glucose-stimulated insulin secretion (GSIS) assay measured insulin secretion in response to 2.8mM (low), 20mM (high) glucose and 20mM glucose + IBMX. Data show average secreted insulin as a percentage of insulin content in the isolated islets from late fetal (n=4) and P22 (n=3) piglets (t-test, *\*\* *P*≤0.01; *** *P*≤0.001*; bars indicate SD). (B) Heat map of DEGs between late fetal and P22 β-cells. The Z-score scale represents log_2_(X+1) transformed transcripts per million (TPM) counts. Red and blue color intensity of the Z-score indicates up-regulation and down-regulation, respectively. (C,E) GO term analysis for biological processes with up-regulated (C) and down-regulated (E) genes between late fetal and P22 β-cells. Adjusted *P* value threshold was 0.1. (D,F) Boxplots displaying normalized TPM (transcripts per million) counts of up-regulated (D) or down-regulated (F) genes between late fetal and P22. Adjusted *P* value * *P*≤0.05; ** *P*≤0.01; *** *P*≤0.001.

To assess the genetic basis of this maturation, we compared transcriptomes from late fetal and P22 β-cells (Fig. 6B). DESeq2 analysis revealed significantly increased expression of 924 genes and decreased expression of 895 genes between late fetal (both E70 and E90) and P22 (Table S8). GO term analysis revealed that up-regulated transcripts were encoded by genes associated with pathways involved in insulin secretion (Fig. 6C and Table S9). Intracellular cAMP and Ca^2+^ levels are crucial regulators of insulin exocytosis (Tengholm and Gylfe, 2017). We observed that transcripts of genes encoding channels activated by cAMP, such as HCN4 and KCNQ1 were up-regulated (Fig. 6D). We also found increased expression of genes regulating cytoplasmic Ca^2+^ concentration or genes involved in Ca^2+^-mediated signaling pathways such as CRACR2A and CAMK2D (Fig. 6D). Phosphodiesterases (PDEs), which induce hydrolysis of cAMP to 5’-AMP, were also up-regulated by stage P22 (Fig. 6C). These findings are consistent with the view that cAMP signaling ‘tone’ may be relatively high in fetal islets, then reduced in the post-natal pig β-cell, a view supported by our quantification of dynamic basal and glucose + IBMX regulation of insulin secretion at these stages (Fig. 6A).

Consistent with our observation that insulin secretion at basal glucose levels declined during fetal to postnatal islet development, we observed that pathways involved in glucose processing such as “glucose catabolic process”, “ATP generation from ADP” and “calcium ion regulated exocytosis” were down-regulated in β-cells (Fig. 6E). Compared to a relative peak in late fetal development, transcripts of genes regulating glycolysis such as *ENO2, ALDOB, GPD1* and *GCK* were down regulated by stage P22 (Fig. 6F). Thus, our data suggests that multiple inter-related signaling pathways regulating insulin exocytosis continue to mature through weaning-age (P22). In summary, our combined developmental, functional and molecular investigations here revealed molecular and genetic factors governing maturation of neonatal β-cells and unveiled conserved mechanisms underlying islet cell development and function in pigs and humans.

## Discussion

Based on the crucial role of pancreatic islets in human diseases and the expanding catalog of differences between human and mouse islet developmental biology or regulation (Arda et al., 2013; Arda et al., 2016; Brissova et al., 2005; Hart and Powers, 2019), there is intense interest in developing complementary experimental systems for investigating islet development and maturation. To address this need, we used an integrated approach to delineate pancreas development in pigs. This multimodal assessment of genetic and developmental phenotypes across a comprehensive range of fetal and post-natal stages of pig pancreas ontogeny and maturation revealed multiple similarities in formation and regulation of pig and human islet β-, α- and δ-cells, including features not observed in mice. Systematic phenotyping of pig pancreas development - particularly gene expression and identification of developmental signaling pathways - revealed unexpected dynamic gene regulation, and evidence of native intra-islet GLP-1 signaling. Moreover, studies in pigs are cost-efficient (see Methods). Our experimental approaches and findings provide a unique roadmap for diabetes and metabolic research, including attempts to direct differentiation and maturation of replacement islet cell types from renewable human cell sources.

Here, we established cell purification strategies based on flow cytometry to investigate developmental genetics in fetal and neonatal pig β-, α- and δ-cells, an approach we and others have successfully used in pancreas from mice and humans (Arda et al., 2016; Blodgett et al., 2015; DiGruccio et al., 2016; Hrvatin et al., 2014a; Qiu et al., 2017). Like in prior studies (Arda et al., 2016; Hrvatin et al., 2014a), we isolated primary β-, α- and δ-cells throughout fetal and postnatal pancreas development using antibodies against hormones specific to these cell types, an approach permitting comprehensive, high-quality stage-specific transcriptome analysis. Similar approaches for primary ductal and acinar cells should provide opportunities in the future to delineate the transcriptome of these important components of the exocrine pancreas. Moreover, discovery of flow cytometry-based methods for purifying live primary endocrine or exocrine cells, with antibodies that recognize cell-surface epitopes, should expand our capacity to investigate other important aspects of pancreatic gene regulation and function, using approaches like ATAC-Seq, promoter capture Hi-C, and Patch-Seq (Arda et al., 2018; Camunas-Soler et al., 2019; Miguel-Escalada et al., 2019).

Transcriptome analysis described here provides evidence of genetic regulatory mechanisms and cell development in pigs that are also conserved in other mammals, including (1) cell type-specific expression of transcription factors (*PDX1, MAFA, ARX, IRX, HHEX*) and functional regulators (*PCSK1, PCSK2, SLC2A2, SLC30A8, GLP1R, GCGR*), (2) transient β-, α- and δ-cell proliferation, peaking in fetal/perinatal stages and regulated by conserved factors like *MKI67, BUB1B* and *NUSAP1* (Georgia and Bhushan, 2004; Meier et al., 2008; Teta et al., 2007), (3) dynamic expression of histone or chromatin regulators like *EZH2* and *DNMT1* (Chakravarthy et al., 2017; Dai et al., 2017) and (4) reduction or elimination of transcripts encoding ‘disallowed’ genes (Lemaire et al., 2017; Pullen et al., 2017; Pullen et al., 2010; Thorrez et al., 2011). These findings from transcriptome analysis also correspond well with observations from our other stage-specific tissue phenotyping (see below). Stage-specific gene expression patterns revealed that pig and human β-cells and α-cells shared characteristic molecular and developmental features not observed in mouse β-cells or α-cells. Like in human islets (Benner et al., 2014), the transcription factor MAFB is expressed in both pig β- and α-cells (Figs. 2,3). Similarly, we found that the transcription factor SIX3 is expressed in pig β-cells in an age-dependent manner, like in human islets (Arda et al., 2016) (Fig. 3). Thus, studies of mechanisms governing cell type- and age-dependent expression of genes like pig *MAFB* and *SIX3* could reveal the basis of β-cell functional maturation. Likewise, we observed features of pig fetal islets not yet noted in humans. *MAFA* expression in pig β-cells increased in fetal stages, and continued through postnatal stages. In humans, β-cell *MAFA* expression begins in fetal stages (Blodgett et al., 2015) and also increases after birth (Arda et al., 2016), but its fetal regulation remains unreported. Our studies also provide a reference transcriptome analysis of developing islet δ-cells, whose recognized role in health and diabetes is growing (Rorsman and Huising, 2018). Together, the resources and methods described here provide evidence that developmental genetic studies of pancreas in pigs could complement similar studies in humans and mice.

In addition to the resemblance of pancreatic gene regulation in pigs and humans (Figs. 2,4), we observed similarities of islet morphogenesis, cell specification, regulation and islet cell allocation. Scattered islet β-, α- and δ-cells coalesced into small clusters at late fetal stages, which contained proliferating cells (Figs. 2,3). At birth, the proportion of α- and δ-cells in pig islets was ∼50%, like that of neonatal human islets, and unlike in mice. In pig pancreas development, a large portion of fetal hormone-expressing islet cells appeared to be contained within small, dispersed clusters of varying cell composition. After birth, these clusters enlarged, and the geometry of β-, α- and δ-cells in islets appeared intermingled, like in human islets (Brissova et al., 2005) (Fig. 3). Studies of insulin secretion further demonstrated that β-cell function matures from fetal to post-natal stages like in humans (Arda et al., 2016) and other mammals. Thus, our study reveals multiple features in pig islet development that are closely conserved in human pancreas, including a subset of phenotypes not observed in rodent islets, suggesting that pigs could be a reliable surrogate animal model to study the islet function of human gene orthologs. This includes studies of dominant forms of diabetes like MODY and NDM, whose genetics have not been reconstituted in mice (Hattersley and Patel, 2017; Maestro et al., 2007). By contrast, dominant monogenic diabetes resulting from mutations in *INS* or the MODY3 gene *HNF1α* has been reconstituted in pigs (Renner et al., 2013; Umeyama et al., 2009). Thus, studies of gene function and pathogenesis in diseases like MODY and NDM could be advanced through investigations in pigs.

Here, we evaluated pancreas phenotypes through a full range of fetal and peri-natal stages. In β-, α- and δ-cells, this ‘granularity’ of phenotyping revealed changes in >6000 genes during development, and identified developmental dynamics not previously noted in studies of human islet gene expression with comparatively limited sampling during developmental stages (Arda et al., 2016; Blodgett et al., 2015; Bramswig et al., 2013), or in one study, a single stage (Ramond et al., 2018). For example, in β-cells, genes involved in glucose processing, including *G6PC2* and *PDK3*, are in the “Up_NC” cluster whereas genes regulating insulin secretion, such as *SLC2A2*, are in the “NC_Up” cluster (Table S6), indicating that mechanisms governing glucose sensing and metabolism, or insulin secretion, are established at different developmental stages in pig β-cells. Moreover, we noted 366 β-cell or α-cell transcripts whose levels changed with a ‘discontinuous’ trajectory in development (Fig. 5 and Table S6). That is, we observed heterogeneous patterns of expression, including early rise then fall (like *DPP4, FEV* in β-cells, and *FOXO1* or *KIT* in α-cells), or initial decline followed by a later rise in transcript levels, including those encoding a subset of disallowed gene products like PDK4 in β-cells (Fig. 5 and Table S6). Thus, our ability to detect dynamic gene expression trajectories in β-cell or α-cells was enhanced by the comprehensiveness of developmental phenotyping afforded in pigs.

Cell- and stage-specific phenotyping here also revealed links between our data to human diabetes risk and islet replacement. Prior studies have noted a significant enrichment of candidate diabetes risk genes among those with dynamic developmental expression (Arda et al., 2016; Perez-Alcantara et al., 2018). Similarly, in our recent work, prioritization of candidate diabetes risk genes with dynamic fetal or post-natal expression led to identification of SIX2, SIX3 and BCL11A as unrecognized regulators of human β-cell development and function (Arda et al., 2016; Peiris et al., 2018). Here, we found significant enrichment of β- and α-cell gene sets involved in specific developmental signaling pathways (Fig. 5). Of these, only a subset, such as mTOR, retinoic acid receptor and TGF-β signaling, have previously been linked to β- and α-cell biology (Blandino-Rosano et al., 2017; Brun et al., 2015; Lin et al., 2009; Yokoi et al., 2016). Thus, our datasets provide a foundation for systematic investigations of *cis-* and *trans-* regulatory elements governing the dynamic expression of diabetes risk genes in the native, physiological setting of pig islet development.

Identifying signals controlling islet development, including detection of signaling modulation, should enhance efforts to control and direct differentiation of functional replacement islet cells from stem cells and other renewable sources. Recent studies have explored use of neonatal pig islet cultures for identifying signals that promote β-cell functional maturation (Hassouna et al., 2018). In their work with stepwise directed differentiation of human stem cells into distinct lineages, Loh and colleagues have demonstrated the value of inhibiting signaling pathways – not merely through signal withdrawal, but by exposure of cells to potent signaling inhibitors – to prevent development along undesired fates or lineages (Ang et al., 2018; Loh et al., 2014). The identification of dynamically regulated genes and pathways, including those with discontinuous trajectories (Up then Down, etc.), in β-cell and α-cell development provides a new roadmap for modulating islet cell replacement efforts. Fetal and neonatal β-cells and α-cells develop in close proximity, and our work likely identifies intercellular signaling pathways (like those regulated by *PCSK1*, *GLP1R* and *DPP4*) that could regulate differentiation of these cells. PCSK1 expression in mouse fetal α-cells was reported previously (Wilson et al., 2002) but active GLP-1 production from fetal islets was not assessed; likewise, to our knowledge, human fetal islet GLP-1 production has not been reported. Recent studies have demonstrated that GLP1R can be stimulated by glucagon, in addition to GLP-1 (Capozzi et al., 2019; Svendsen et al., 2018; Zhu et al., 2019); however, it remains unknown if all signaling downstream of GLP1R is activated by glucagon, or if islet cells are the sole target of GLP-1 signaling in the fetal pancreas. Further studies of endogenous GLP-1 signaling in pig fetal islet development could also clarify the emerging concept of intra-islet GLP-1 signaling in metabolically-stressed adult islets (Capozzi et al., 2019; Drucker, 2013). Thus, our findings identify multiple pathways associated with fetal and post-natal β-cell maturation or function (Table S9), including many not directly modulated in prior studies (Nair et al., 2019; Pagliuca et al., 2014; Rezania et al., 2014; Russ et al., 2015). In summary, data, conceptual advances and resources detailed here provide an unprecedented molecular, cellular and developmental framework for investigating pancreas development that could inform our understanding of diabetes and obesity risk, and advance efforts to produce replacement human islets for diabetes.

## Materials and methods

### Animals and pancreas harvest

Pigs used in this study were outbred and used in accordance to protocols approved by the Institutional Animal Care and Use Committees at University of California Davis and Stanford University. For collection of fetal pig pancreata, pregnancies were confirmed via ultrasound 30 days after mating and pregnant gilts were humanely euthanized at 40, 70, or 90 days. The reproductive tracts were removed after mid-ventral laparotomy, cooled on ice, and fetuses were collected. Fetuses remained on ice until dissection. Pancreas was identified according to anatomical location and harvested under a stereoscopic microscope. Neonatal pancreata were harvested from neonatal piglets at 8 and 22 days of age, while adult pancreata were collected from pigs 6 months of age or older. Briefly, after euthanasia, a midline incision was performed to provide access to the peritoneum and thoracic cavity. The pancreas was rapidly chilled via either perfusion of sterile ice-cold saline solution into the descending aorta or by pouring the same solution directly into the peritoneal cavity and then surgically removed. Harvested pancreatic tissue was maintained at a temperature of 2-10°C in saline solution until further downstream processing.

### Generation of pancreatic single cell suspensions

Morphological differences of the pancreas between developmental stages necessitated customized protocols to generate single cell suspensions suitable for downstream assays. Embryonic day 40 pancreata were diced finely with a razor blade and transferred into a 4°C solution of Collagenase D (Millipore-Sigma, Cat# C9263) at a concentration of 1 mg/ml in HBSS, and digested to single cells overnight at 4°C. Embryonic day 70 and 90 pancreata were finely diced with a razor blade and digested for 5-8 minutes at 37°C using the same Collagenase D solution above. The reaction was stopped with the addition of equal volume HBSS containing 10% porcine serum (Millipore-Sigma, Cat# P9783). If necessary, an additional digestion using TrypLE (Gibco, Cat#12605-010) for 3 minutes at 37°C was used to fully disperse any remaining cell clusters. Post-natal pancreata were digested following published methods (Lamb et al., 2014). (Logistical and technical challenges related to the mass of adult pigs precluded isolation of single cell islet suspensions of sufficient quality suitable for molecular analysis of adult pig islets.) Briefly, pancreata were trimmed to remove lymphatic, connective, and fat tissues, and chopped into 1-5 mm pieces. The tissue was digested using 2 mg/ml Collagenase D in HBSS at 37°C for 12-18 minutes. 1.5 volumes of HBSS with 10% porcine serum and 0.5% Antibiotic/Antimycotic solution was added to stop digestion. Neonatal Islet Cell clusters (NICCs) of the desired size were obtained after straining digest through a 500 μm mesh (PluriSelect, Cat# 43-50500-03) to remove large undigested tissue, washed, and allowed to gravity settle two times. To generate single cell suspensions, some NICCs were further dispersed with TrypLE for 3 minutes at 37°C. Remaining NICCs were spun down and resuspended in RPMI 1640 (Gibco, Cat# 11879-020) containing 5 mM glucose, 10% porcine serum, 10 mM HEPES (Caisson Labs, HOL06-6X100ML) and 1% Pen/Strep (Life Technologies, Cat# 15140-122) and placed in a humidified, 5% CO_2_ tissue culture incubator.

The cost basis of pancreas procurement and islet isolation from pigs is currently lower than for humans. In 2019, reimbursement for a single human cadaveric donor pancreas is between $4,500 and $7,000 (US dollars). For human islets, investigators are charged $0.12 per islet equivalent using the NIH-supported Integrated Islet Distribution Program (https://iidp.coh.org). By contrast, the cost of a P22 pig pancreas is approximately $500, and after processing, a P22 pancreas yields approximately 90- to 100,000 NICCs. The total cost per pig NICC is <$0.008.

### Cell flow cytometry and purification

Single cell suspensions were washed twice with cold PBS and filtered with a 70 µm filter (BD Falcon, Cat# 352350). Prior to fixation, cells were stained with LIVE/DEAD fixable Aqua (ThermoFisher, Cat# L34976) or LIVE/DEAD fixable Near-IR (ThermoFisher, Cat#L10119) dead cell stains according to manufacturer’s protocol. Cells were fixed with 4% PFA in PBS (Electron Microscopy Sciences) for 10 minutes at room temperature. After fixation, cells were permeabilized with Permeabilization Buffer: 1X PBS containing permeabilization solution (BioLegend, Cat# 421002), 0.2% BSA (Sigma-Aldrich, Cat#A9576), mouse and rat IgG (Jackson Labs, Cat# 015-000-003 and 012-000-003), and 1:40 Ribolock RNase Inhibitor (Fisher Scientific, Cat# FEREO0384), for 30 minutes at 4°C. Cells were pelleted and resuspended with Wash Buffer: 1X PBS containing permeabilization solution, 0.2% BSA, and 1:100 Ribolock. Cells were pelleted again, supernatant removed, and stained with Alexa Fluor 488 labeled anti-insulin (R&D Systems, IC1417G-100UG), Alexa Fluor 647 labeled anti-glucagon (Santa Cruz, sc-57171 AF647) and PE labeled anti-somatostatin (Santa Cruz, sc-55565 PE) antibodies in Staining Buffer: 1X PBS containing permeabilization solution, 1% BSA, and 1:40 Ribolock, for 30 minutes at 4°C. Cells were washed three times in Wash Buffer and resuspended in Sort Buffer: 1X PBS containing 0.5% BSA, and 1∶20 Ribolock. Labeled cells were sorted on a special-order, 5-laser FACS Aria II using FACSDiva software (BD Biosciences). After excluding doublets and dead cells, desired cell populations were identified using fluorescence minus one (FMO) controls. Cells were sorted using the purity mask in chilled low retention tubes containing 50 μl of Sorting Buffer. We attempted to isolate cells producing more than one islet hormone (insulin, glucagon or somatostatin) simultaneously, which have been reported in both human and pig fetal pancreas (Lukinius et al., 1992; Riopel et al., 2014). In contrast to these studies, however, the relative frequency of these cells after FACS was quite low (<0.001% of live cells) and precluded downstream analysis. Similarly, the relative paucity of δ cells in the early fetal porcine pancreas has proven a significant hurdle to collection of sufficient numbers for downstream analysis. Post-hoc visualization of recorded data was performed using FlowJo vX software (FlowJo, LLC).

### Total RNA extraction and quantitative Reverse Transcription PCR (qRT-PCR)

Total RNA was extracted from the sorted cells as described (Hrvatin et al., 2014a) and concentrated using the RNA Clean & Concentrator-5 kit (Zymo Research, Cat# R1013). Complementary DNA (cDNA) was generated using the Maxima First Strand cDNA Synthesis kit (Thermo Scientific, Cat# K1642) according to the manufacturer’s instructions. Purity of sorted populations was confirmed via Quantitative PCR using an Applied Biosystems 7500 Real-Time PCR System. The following TaqMan probes were used: INS (Ss03386682_u1), GCG (Ss03384069_u1), SST (Ss03391856_m1), AMY2 (Ss03394345_m1), ACTB (Ss03376081_u1), KRT19 (Forward: 5’-GAAGAGCTGGCCTACCTG-3’, Reverse: 5’-ATA CTGGCTTCTCATGTCGC-3’), SIX2 (Forward: 5’-TCAAGGAAAAGAGTCGCAGC-3’, Reverse: 5’-TGAACCAGTTGCTGACCTG-3’) and SIX3 (Forward: 5’-AACAAGCACGAGTCGATCC-3’, Reverse: 5’-CCACAAGTTCACCAAGGAGT-3’). Relative mRNA abundance was calculated by the Comparative Ct (ΔΔCt) relative quantitation method.

### RNA-Seq Library preparation

Multiplexed libraries were prepared using the SMARTer Stranded Total RNA-Seq Kit v2 - Pico Input Mammalian (TakaraBio, Cat# 634413) following manufacturer’s instructions. Library fragments of approximately 350 bp were obtained, and the quality was assessed using a 2100 Bioanalyzer (Agilent). Barcoded libraries were multiplexed and sequenced on an Illumina NextSeq sequencer.

### Bioinformatic and statistical analysis of RNA-Seq datasets

RNA-Seq libraries were sequenced as 75 bp pair-end reads to a depth of 30 to 60 million reads. To estimate transcriptome abundance, Salmon algorithm (v0.11.3) was used (Patro et al., 2017). An index was built on the Sus Scrofa reference transcriptome (Sscrofa 11.1, version 96) (Zerbino et al., 2018) using Salmon with parameters “salmon index --keepDuplicates -t transcripts -i transcripts_index --type quasi -k 31”. The reads were aligned to the index and quantified using Salmon with parameters “salmon quant -i $salmon_index -l A \-1 $Input1 \-2 $Input2 \ p 8 -o quants/${Input1}_quant \ --gcBias –seqBias”. After getting the transcript counts, differentially expressed gene analysis was performed using the DEseq2 analysis pipeline (Love et al., 2014; Soneson et al., 2015). Differentially expressed genes (DEGs) were obtained by comparisons of any two sequential stages (i.e. Early fetal versus Late fetal and Late fetal versus Neonatal) in a cell type specific manner. Genes were considered differentially expressed if the fold change between samples was greater than or equal to 1.5 with setting of Benjamini-Hochberg (BH) false discovery rate (FDR) adjusted P value of <0.05 in the package. Hierarchical clustering of samples were done with the cor() functions of R. Principal component analysis (PCA) was performed with the function *plotPCA* that comes with DEseq2 package. Visualization of RNA-seq data examined in this study were produced with pheatmap 1.0.12 (Kolde, 2012) and ggplot2 (Wickham, 2009). GO term analysis was carried out using the R package clusterProfiler as described (Yu et al., 2012) with a Benjamini-Hochberg adjusted P value of <0.1.

### Comparison to extant human and mouse β-cell transcriptomes

Human (Arda et al., 2016; Blodgett et al., 2015) (GEO accession numbers GSE67543 and GSE79469) and mouse (Qiu et al., 2017) (GEO accession number GSE87375) datasets were obtained from the NCBI Gene Expression Omnibus (GEO). Indices for human and mouse libraries were built from reference transcriptomes GRCh38 cDNA and GRCm38 cDNA, respectively (Homo sapiens, version 96; Mus musculus, version 96; (Zerbino et al., 2018)). Pig and mouse Ensembl IDs were converted to their corresponding human Ensembl ID using predicted ortholog tables across these species obtained from Ensembl BioMart (Zerbino et al., 2018). Sample-wise hierarchical clustering analysis (Fig. 4A) was performed using dist() function of R.

### Identifying genes with dynamic expression

To create lists of dynamically expressed genes, DEGs from comparisons between early fetal and late fetal stages and between late fetal and neonatal stage were intersected, which created lists of genes showing patterns of gene expression with up, down or no change directions as development proceeds in islets development. Gene TPM values shifted by +1, log based 2 transformed were averaged across biological replicates and subsequently standardized to their Z-scores.

### Identification of developmentally regulated MODY and NDM genes

We queried 33 genes previously linked to MODY and/or NDM (Velayos et al., 2017; Yang and Chan, 2016), and identified 19 dynamically expressed genes during the transition from fetal to neonatal stages in pig β-, α- or δ-cells. Among these 19 candidates, we then used prior data sets (Blodgett et al., 2015) to identify a subset that also have dynamic expression in human islet β- or α-cells – a lack of available human δ-cell datasets precludes this comparison for δ-cells.

### Immunohistochemistry

Biopsies of harvested pancreata were fixed in 4% paraformaldehyde (w/v) in PBS at 4°C overnight, washed twice in PBS, and cryoprotected via sucrose saturation. The tissues were imbedded in Tissue-Tek OCT compound (VWR, Cat# 25608-930) and stored at -80°C until sectioning. 10μm thick sections were collected on ThermoFisher Superfrost Plus slides. For staining, frozen slides were thawed at room temperature for 30 min and washed with PBS. Antigen retrieval was performed by boiling slides immersed in Target Retrieval Solution (DAKO, S169984-2) for 30 minutes, as needed. Subsequently, sections were incubated with streptavidin/biotin blocking solution (Vector Labs, SP-2002) according to manufacturer’s instruction. Then, sections were incubated in permeabilization/blocking buffer (5% normal donkey serum, 0.3% Triton X-100 in PBS) at room temperature for 1 hour followed by overnight incubation at 4°C with primary antibodies diluted in Antibody dilution buffer (1% bovine serum albumin, 0.3% Triton X-100 in PBS). Slides were washed and stained with appropriate secondary antibodies for 1 hour at room temperature. Primary antibodies and dilutions used are as follows: guinea pig anti-insulin (DAKO, A0564, 1:1000), mouse anti-glucagon (Sigma-Aldrich, G2654, 1:1000), rabbit anti-SIX3 (LifeSpan Biosciences, LS-B9336, 1:100), goat anti-PDX1 (R&D systems, AF2419, 1:200), mouse anti-human NKX6.1 (R&D systems, AF5857, 1:200,) rabbit anti-MAFB (Bethyl, IHC-00351, 1:200), rabbit anti-MAFA (AVIVA systems biology, ARP47760_P050, 1:200), goat anti-somatostatin (Santa Cruz biotechnology, 1:500), rabbit anti-Chromogranin A (Immunostar, 20085, 1:500), rat anti-E-cadherin (Life Technologies, 13-1900 1:500). Secondary antibodies used are donkey anti-isotype conjugated with Alexa Fluor 488/555/647 (1:500; Molecular Probes). After a final wash with PBS, slides were preserved with a mounting medium containing DAPI (Vector labs, H-1200) and coverslipped. Image acquisition was performed on a Leica SP8 confocal microscope. Islet cells were quantified by counting insulin positive β-cells, glucagon positive α-cells, and somatostatin positive δ-cells using pancreas sections from different stages.

### Morphometric Analysis

Tissue sections were prepared as above. Single-channel images were captured using a Zeiss Axio Imager M2 microscope using a 20X objective with the AxioVision version 4.8 software. Respective channel images were then imported into the Image-Pro Plus version 5 software (Media Cybernetics, Inc.). Quantification of hormone^+^ cells was performed by using the count function within the Measure menu. The number of individual hormone^+^ cells were then summated for each developmental stage, and the fractional percentage comprising each of the three cell types was calculated. To determine the percentage of β-, α-, and δ-cells which were undergoing cell division at each developmental stage, slides were stained for a combination of insulin, glucagon, and Ki67 or insulin, somatostatin, and Ki67. Single-channel images were collected as above and then merged into a multi-channel image using the Image-Pro Plus software. Cells which were Ki67^+^ and hormone^+^, as well as the total number of hormone^+^ cells, were then counted manually.

### In vitro insulin secretion assay

Functional maturity of NICCs isolated from late fetal or 22-day old piglets was assessed by in vitro glucose stimulated insulin secretion (GSIS) assay performed after 5 days of culture. In brief, 100 NICCs were washed and incubated at 2.8 mM glucose for 1 hour. After the initial preincubation, clusters were serially incubated in media above containing 2.8 mM, 20 mM, and 20 mM plus IMBX glucose concentrations for 1 hour each. Finally, clusters were lysed in acid-ethanol solution to extract the total insulin content. Insulin from supernatants and the NICC lysates was quantified using the pig insulin ELISA kit (Mercodia, 10-12000-01). Secreted insulin is presented as a percent of total insulin detected in lysate. All steps were performed in triplicate at 37°C using RPMI 1640 supplemented with 2% porcine serum adjusted to the respective glucose concentrations.

### In vitro active GLP-1 assay

Islet cell clusters were handpicked immediately after isolation, washed twice in ice cold PBS, then resuspended in 4°C lysis buffer containing DPP4 inhibitor (EMD Millipore, DPP4). Clusters were sonicated at 4°C and immediately stored at -80°C. Active GLP-1 (7-36) was measured using the Mouse / Rat GLP-1 Active (7-36) ELISA Kit (Eagle Biosciences, GP121-K01).

### Statistical analysis

For qRT-PCR, GSIS and quantification of immunohistology experiments, the number of biological or technical replicates (n), measure of central tendency (e.g. average), standard deviation (SD), and statistical analysis is detailed in the Figure legend. Graphs and statistical analysis were produced and performed by using GraphPad Prism (version 8) software.

## Acknowledgements

We thank past and current members of the Kim group for advice and encouragement, Dr. Sam Baker (Stanford, Dept. of Comparative Medicine, Veterinary Service Center), Dr. Jing Wang (Stanford Diabetes Research Center and Stanford Islet Research Core) and Michael Alexander (U.C. Irvine) for assistance with pig pancreas work, Serena Whitener for help developing the pig pancreas gene expression browser, Dr. Kyle Loh for stimulating discussions, and Drs. Dan Lu and Ramesh Nair for help on bioinformatics.

## Data Availability

The data discussed in this publication have been deposited in NCBI’s Gene Expression Omnibus (Edgar et al., 2002) and are accessible through GEO Series accession number GSE143889 (https://www.ncbi.nlm.nih.gov/geo/query/acc.cgi?acc=GSE143889).

## Author contributions

S.K., R.L.W. and S.K.K. designed the study; S.K, R.L.W, X.G., C.A.C., J.Y.L., I.P., R.J.B., and K.T. performed the experiments; J.C., S.R.Q., J.R.T.L., R.B., and P.J.R. shared materials; S.K., R.L.W., H.P., and S.K.K analyzed the data and wrote the manuscript.

