## Supplementary Figures for "Molecular and genetic regulation of pig pancreatic islet cell development"

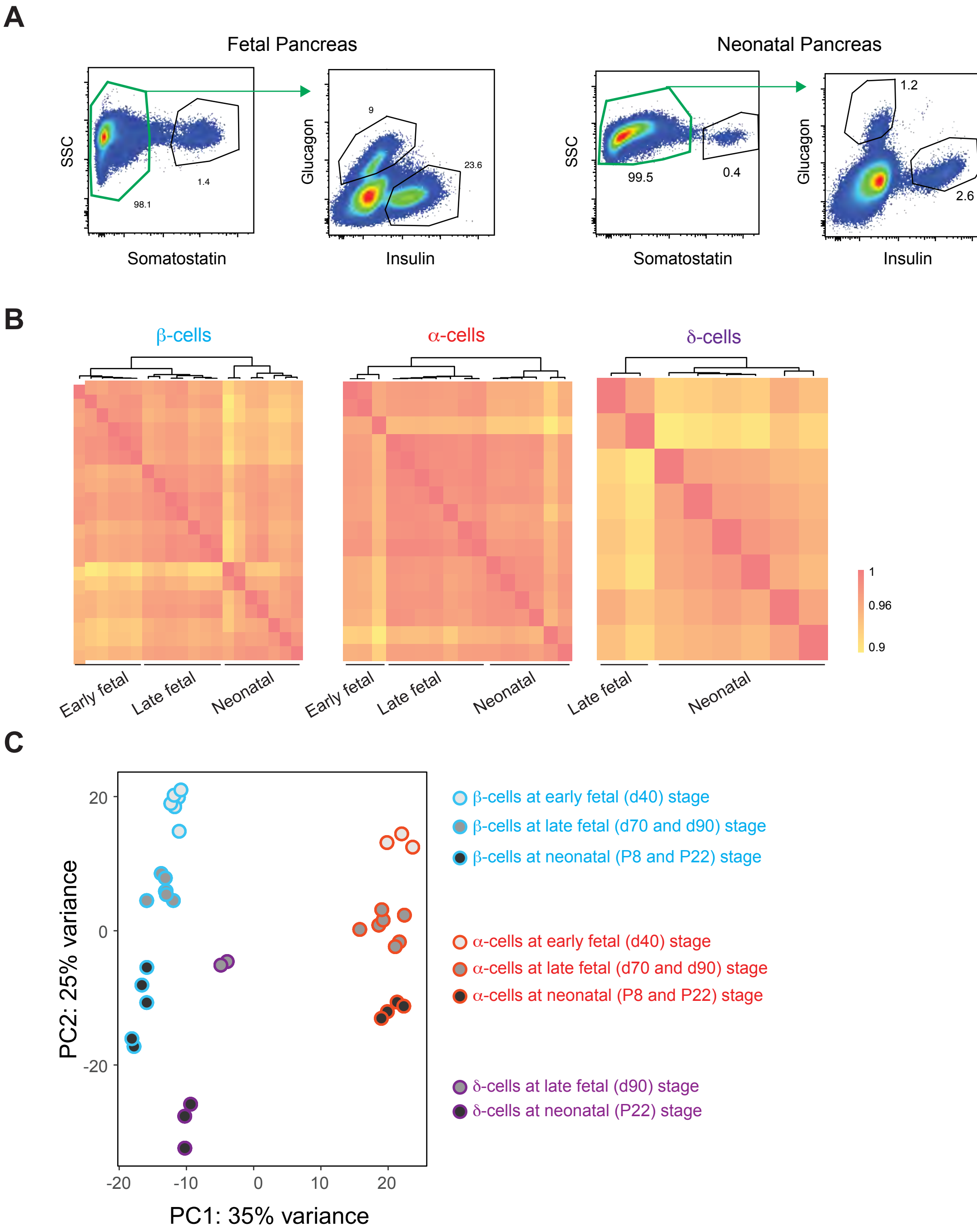


Fig. S1. (A) FACS plots demonstrating distinct populations of Insulin (INS) positive β-, glucagon (GCG) positive α- and somatostatin (SST) positive δ-cells among dispersed fetal and neonatal pancreatic cells. (B) Pearson correlation matrix of all RNA-seq samples used in this study demonstrating distinct stage specific gene expression patterns in β-, α-, and δ-cells (C) Principal Component Analysis indicating the RNA-Seq libraries cluster according to cell type in principle component 1, followed by developmental stage in principle component 2

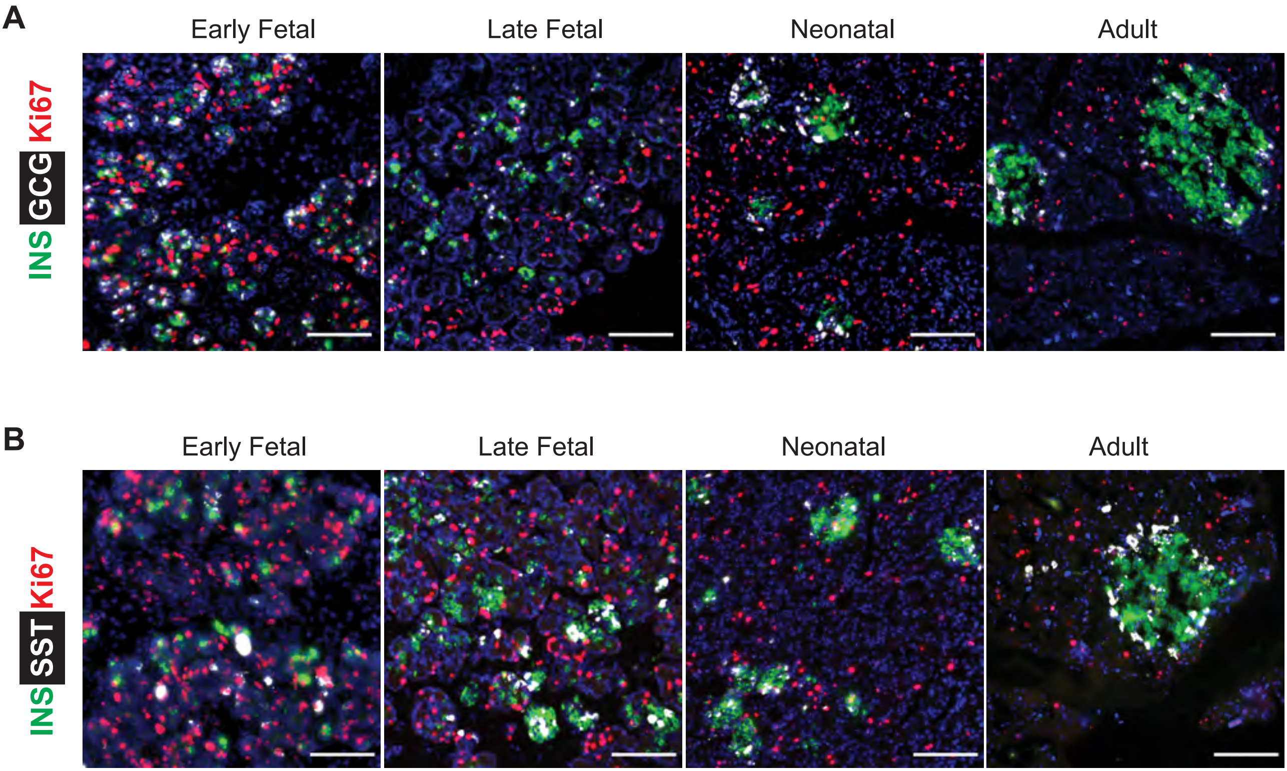


Fig. S2. Representative immunofluorescence images showing (A) Insulin and Glucagon or (B) Insulin and Somatostatin co-stained with Ki67 (Scale bars, 100 μm)

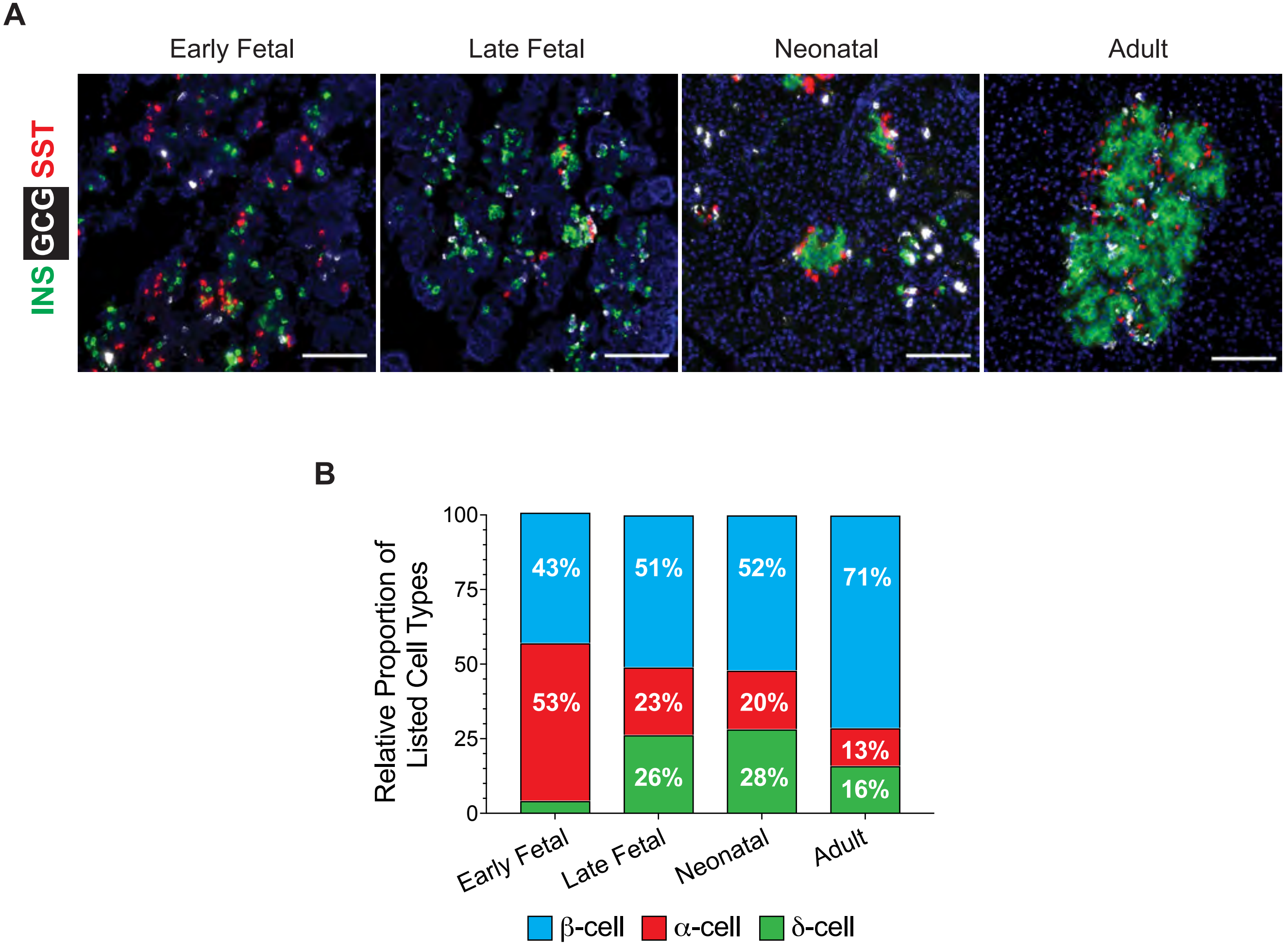


Fig. S3. Representative immunofluorescence images (A) and (B) their quantification graphs showing relative proportion of INS+, GCG+ and SST+ cells in each developmental stage (n = 40 images per group, from 3 pigs per group, Scale bars, 100 μm)

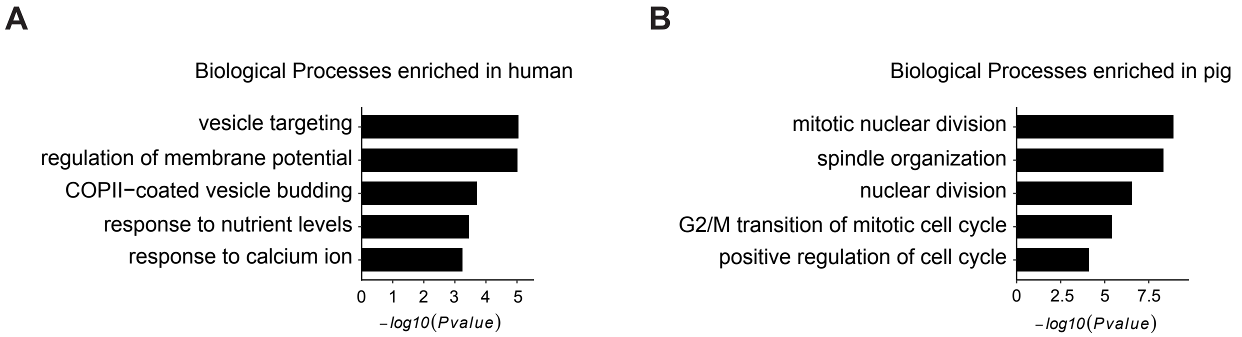


Fig. S4. Enriched GO terms with genes enriched in (A) human and (B) pig β-cells at neonatal stage. Adjusted p value threshold was 0.05.

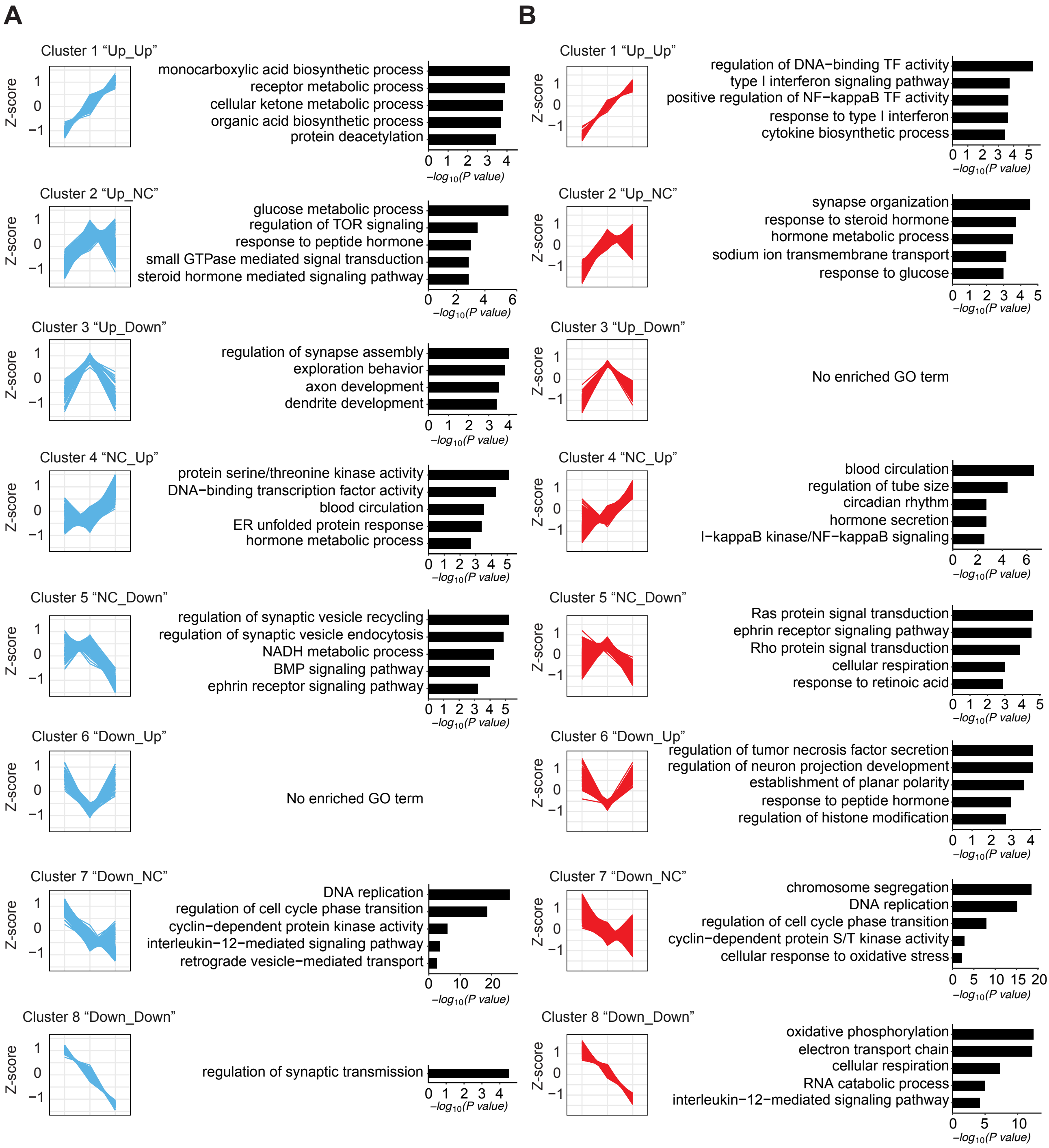


Fig. S5. (A, B) Visualization of gene expression change with z score representing log2(X+1) transformed TPM counts of genes in 8 clusters. Genes were clustered based on expression changes from early fetal stage to neonatal day 22 in β- (A) and α- (B) cells. Enriched GO terms with genes in each cluster are shown to the right. Adjusted p value threshold was 0.1.

**Supplementary Tables**

**Table S1.** List of RNA-Seq libraries sequenced in this study

**Table S2**. Differentially expressed genes in pig β-, α-, and δ-cells between two developmental stages (Early fetal vs Late fetal, Late fetal vs Neonatal)

**Table S3.** TPM counts of genes in each RNA-seq library

**Table S4.** Genes which are developmentally regulated in human and pig islet cells

**Table S5.** Differentially expressed genes and GO term analysis comparing neonatal human and pig β-cells.

**Table S6.** Genes with dynamic expression in pig β- and α-cells from early fetal to neonatal stages.

**Table S7.** MODY and NDM genes which are developmentally regulated in human and pig islet cells

**Table S8.** Up- and down-regulated genes in pig β-cells between late fetal and P22.

**Table S9**. GO term analysis in pig β-cells between late fetal and P22.
